# E3 ligase Praja1 mediates ubiquitination and degradation of microtubule-associated protein Tau

**DOI:** 10.1101/2024.06.10.598176

**Authors:** Shiho Aoki, Kotaro Kawasaki, Kazuki Imadegawa, Mizuho Oishi, Toru Asahi, Wataru Onodera

## Abstract

RING-H2 type E3 ligase Praja family is composed of Praja1 and Praja2, which promote the degradation of substrates through the ubiquitin-proteasome system. Both paralogs contribute to neuronal maturation and differentiation, indicating a significant role in the nervous system. Aggregation-prone proteins associated with neurodegenerative diseases, including TDP-43 and α-synuclein, are degraded and/or suppressed by Praja1. Furthermore, the expression level of the *MAPT* gene, which is frequently mutated in Alzheimer’s, is regulated by Praja2. While the Praja family has been shown to recognize various aggregation-prone proteins as substrates, it has not been determined whether Tau, a key protein that aggregates in tauopathies, is also recognized by Praja proteins. In this study, we show that Praja1, but not Praja2, recognizes Tau as a candidate substrate. We observed that Tau expression in human neuroblastoma SH-SY5Y cells decreased depending on the E3 ligase activity of Praja1. Furthermore, Praja1 polyubiquitinated and interacted with Tau, indicating that it is a target substrate. Next, by combining ancestral sequence reconstruction and mutational analysis, we revealed that the Praja1-Tau interaction began via deletion of the N- and C-terminal regions of Praja1, occurring just after the duplication of the Praja family in the common ancestor of placentals. Lastly, to test whether this interaction is disrupted under pathological conditions, P301L Tau was introduced, resulting in a degradation similar to that of wild-type Tau. These results reveal an unidentified mechanism of Tau proteostasis by Praja1 and may provide insight into the pathogenesis of neurodegenerative diseases, including tauopathy.

## 1. Introduction

The Praja family has been identified as a RING-H2 type E3 ligase that functions in the ubiquitin-proteasome system to maintain intracellular proteostasis through substrate degradation [1-3]. In humans, this family consists of two paralogs, Praja1 and Praja2. Our previous study showed that these paralogs duplicated during evolution to placentals and that Praja1, in particular, may have functionally differentiated from Praja2 due to its increase in the evolutionary rate immediately after duplication [2].

The Praja family is expressed ubiquitously, with prevalence in brain regions, such as the cerebellum and frontal cortex [1]. The physiological function of Praja is thought to be associated with neuronal morphology, and positive and negative regulation of neurite outgrowth has been reported [4,5]. Studies indicate that Praja1 and Praja2 are associated with neuronal diseases. For example, TDP-43, a major component of cytoplasmic aggregates present in amyotrophic lateral sclerosis (ALS) and frontotemporal lobar degeneration (FTLD), is a substrate of Praja1, and their interaction results in resulting in the inhibition of its phosphorylation and aggregation [6,7]. Additionally, aggregation of proteins, including FUS, SOD1, α-synuclein, ataxin-3, and huntingtin polyglutamine proteins, is inhibited by Praja1 [8,9]. The expression levels of Praja2 are repressed in Alzheimer’s disease (AD), which is characterized by amyloid-beta fibrosis and microtubule-associated protein tau (Tau) aggregate formation [10].

Tau, encoded by the *MAPT* gene, is mainly involved in axonal transport and synaptic plasticity and is expressed in the central and peripheral nervous systems [11]. In neurons, it is primarily localized in the cytoplasm and regulates neuronal morphology by binding to and stabilizing microtubules [12,13]. Alternative splicing of *MAPT* transcripts generates six major isoforms of human Tau, and an imbalance between isoform expression or misfolding is observed in several neurodegenerative diseases, such as AD, Parkinson’s disease (PD), and frontotemporal dementia (FTDP), all termed tauopathies [14-16].

Currently, it is unclear whether the Praja family recognizes Tau as a substrate, along with other aggregation-prone proteins. Clarifying this protein interaction may provide novel insights into the mechanisms of maintaining tau protein homeostasis disrupted in tauopathy. Here, we found reduced Tau protein levels when Praja1, but not Praja2, was overexpressed in human neuroblastoma SH-SY5Y cells. Pharmacological and mutagenesis experiments indicated that Tau is recognized by Praja1 and is ubiquitinated and degraded as a candidate substrate. For its recognition, Praja1 requires a deletion mutation that occurred during its evolution in the common ancestor of placentals. The artificial deletion of the region from Praja2 enabled it to recognize Tau. Finally, we showed that the FTDP Tau mutation P301L, which often disrupts the interaction between the physiological binding partners of Tau, is recognized and degraded by Praja1. Our study provides a potential mechanism of Tau proteostasis regulated by the E3 ligase Praja1, which may be particularly valuable in diseases such as tauopathy, in which Tau proteostasis is disrupted.

## 2. Results

### 2.1 Praja1 regulates Tau stability in human neuroblastoma SH-SY5Y cells

As the Praja family regulates neurodegenerative disease-associated genes and their products, we hypothesized that they could potentially interact with Tau, which is also closely related to neurodegenerative diseases. To verify this, Tau was expressed together with Praja family proteins in human neuroblastoma SH-SY5Y cells, and the protein levels were biochemically analyzed. Tag-free 2N4R Tau and N-terminally FLAG-tagged Praja1/Praja2 were used. Co-expression resulted in a statistically significant reduction in Tau protein levels for Praja1 but not for Praja2 (Fig. 1A, B). Notably, both the Tau and Praja family proteins were expressed in the cytoplasm (Fig. 1C). However, endogenous Tau levels were not negatively regulated by Praja1 (Supporting Information 1); there was a difference in the molecular weights of endogenous and transfected Tau, and this may be explained by Tau isoform-dependent regulation by Praja1.

**Figure 1.**
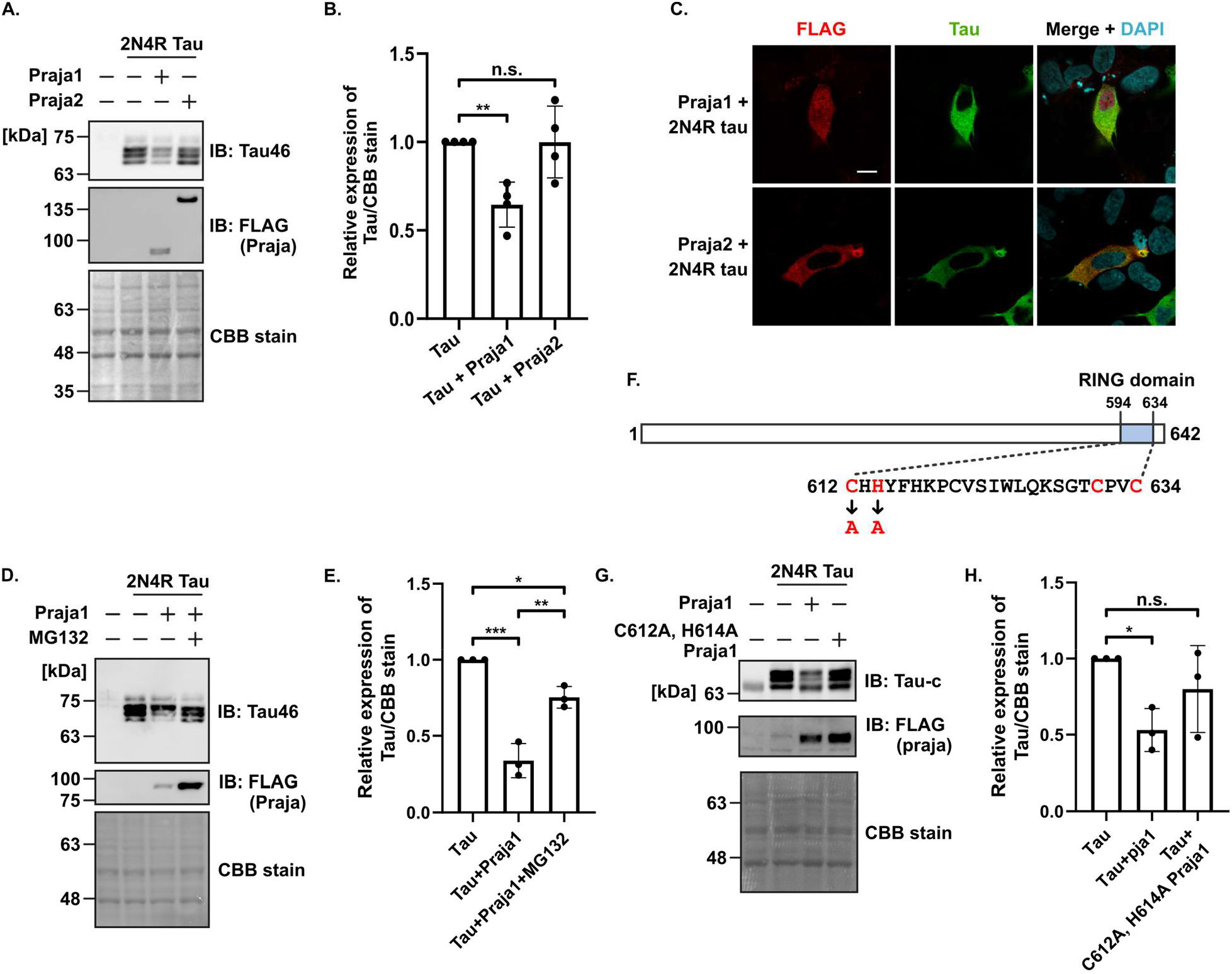
Praja1 negatively regulates Tau in an E3 ligase activity-dependent manner in SH-SY5Y cells. A. Western blot for FLAG-tagged Praja1/Praja2 and Tau co-expressed in SH-SY5Y cells. B. Quantification of the results of A. C. Immunostaining of FLAG-Praja and Tau in SH-SY5Y cells transfected with FLAG-tagged Praja1/Praja2 and Tau alone or co-transfected. DAPI was used for nuclear staining. Scale bar: 10 µm. D. Western blotting for Tau in SH-SY5Y cells treated with or without MG132 for 6 h and co-expressed with Praja1. E. Quantification of the results of D. F. Graphical representation of the RING finger domain of Praja1 (blue) and the zinc finger motif of the RING finger domain (red). G. Co-expression of Praja1 mutated in the zinc finger motif of the RING finger domain (C612A, H614A Praja1) and its western blot analysis. H. Quantitative results of the results of D. For statistical analyses, mean values were subjected to multiple testing by ANOVA with post hoc Dunnett test (*p < 0.10, **p < 0.05, ***p < 0.01).

Next, we examined whether the reduction in Tau was dependent on the E3 ligase activity of Praja1. First, the cells were treated with MG132 for 6 h to inhibit the proteasomal degradation. MG132 treatment restored Tau protein levels, which were otherwise reduced by Praja1 (Fig. 1D, E). Furthermore, two mutations (C612A and H614A) were introduced into the RING domain of Praja1, which is essential for its E3 ligase activity (Fig. 1F) [17]. Upon expression of C612A and H614A Praja1, no decrease in Tau was observed compared to the wild-type Praja1 (Fig. 1G, H). These results suggest that Praja1 negatively regulates Tau stability in SH-SY5Y cells in an E3 ligase activity-dependent manner.

### 2.2 Interaction between Praja1 and Tau

Based on the above results, we hypothesized that Tau is a substrate of Praja1. To confirm this, the direct interaction between Praja1 and Tau was examined using a pull-down assay. First, N-terminal His-tagged Praja1 was purified from *E. coli* and confirmed by CBB staining and western blotting (Supporting Information 2A-C). Unexpectedly, Tau seemed to not interact directly with His-Praja1 under our experimental conditions indicating that the interaction between Praja1 and Tau may be weak and transient (Supporting Information 2D).

Subsequently, Venus-based Bimolecular fluorescence complementation (BiFC) assay was utilized to visualize interaction of the two proteins in live cells [18]. We generated fusion proteins by attaching the N-terminal (VN155) and C-terminal (VC155) fragments of Venus to either the N-terminus or C-terminus of Tau, creating VC155-Tau and Tau-VC155 constructs, respectively. Additionally, we generated VN155-Praja1 by fusing VN155 to the N-terminus of Praja1. Immunofluorescence analysis confirmed that these BiFC constructs exhibited subcellular localization comparable to that of FLAG-Praja1 and 2N4R-Tau (Supporting Information 3). Co-expression of VN155-Praja1 with VC155-Tau yielded cytoplasmic fluorescence, indicating an interaction between these proteins (Fig. 2A). Conversely, co-expression of VN155-Praja1 with Tau-VC155 failed to generate fluorescence (data not shown). We next investigated whether Praja1-dependent polyubiquitination of Tau occurs. Co-immunoprecipitation using a Tau antibody showed that Praja1 promotes Tau polyubiquitination under MG132 (Fig. 2B). Furthermore, co-localization of Praja1 and Tau was observed in the cytoplasm upon treatment with MG132 (Fig. 2C). Taken together, we conclude that Tau is a possible substrate of Praja1.

**Figure 2.**
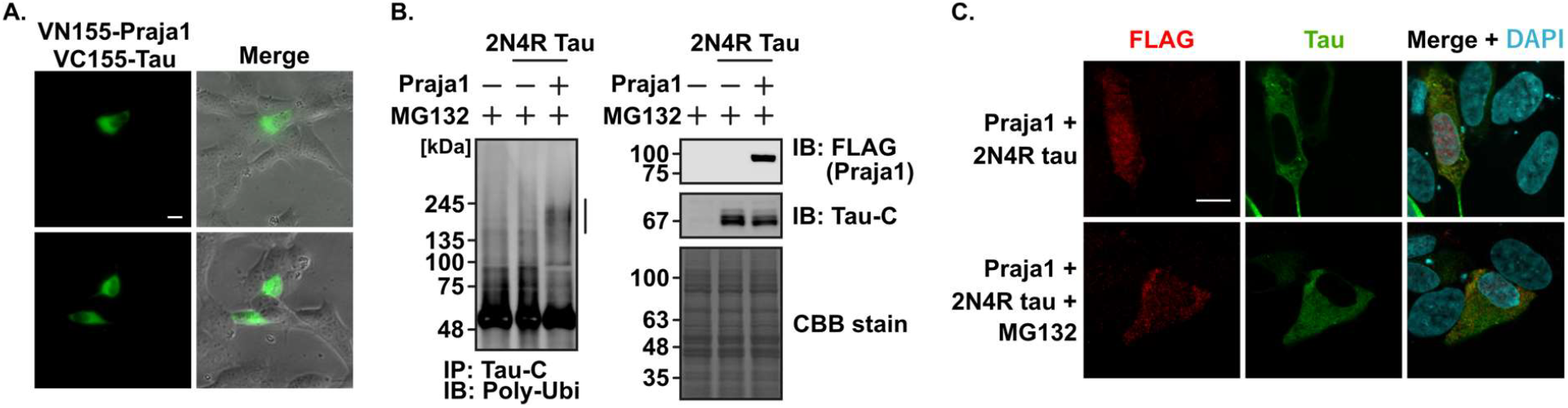
Praja1 may interact and ubiquitinate Tau as a substrate. A. VN155-Praja1 and VC155-Tau co-expressed in the SH-SY5Y cells and Venus-based BiFC fluorescence was observed by fluorescence microscopy. B. FLAG-Praja1 and Tau co-expressed in the SH-SY5Y cells treated with MG132 for 6h. Immunoprecipitation was performed with Tau-C antibody, and multi-ubiquitin was detected by Western blotting. Vertical line shows potential ubiquitinated Tau. C. Immunostaining for FLAG-Praja and Tau in SH-SY5Y cells transfected with both FLAG-Praja1 and Tau, treated with MG132 for 6 h. DAPI was applied for nuclear staining. Scale bar: 10 µm.

### 2.3 Praja1-specific evolutionary deletion is required for Tau recognition

Next, we attempted to understand the evolutionary mechanism underlying Tau and Praja1 interaction. To address this, we reconstructed the common ancestral sequences of Praja1 and Praja2 (Anc-Praja), which are the common ancestors of the placenta (Fig. 3A), using graphical representations of ancestral sequence predictions (GRASP) [19].

**Figure 3.**
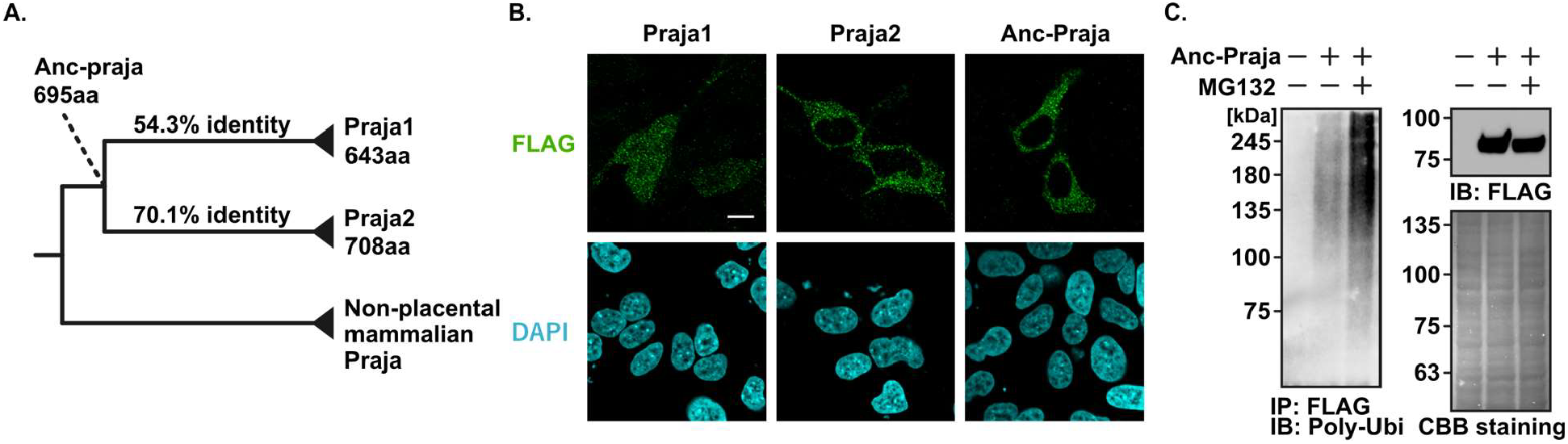
Resurrected ancestral Praja possess E3 ligase activity. A. Phylogenetic position of ancestral Praja. B. Immunostaining of FLAG-Praja and Tau transfected together into SH-SY5Y cells. DAPI was used for nuclear staining. Scale bar: 10 µm. C. FLAG-Anc-Praja and Tau co-transfected SH-SY5Y cells were treated with MG132 for 6 h. Immunoprecipitation was performed using FLAG antibody, and multi-ubiquitin was detected by Western blotting.

Anc-Praja showed a similarity of 54.3% and 70.1% to Praja1 and Praja2, respectively, with lengths of Praja1 (643aa) < Anc-Praja (695aa) < Praja2 (708 aa). Sequence conservation between Anc-Praja, Praja1, and Praja2 was relatively high in the N- and C-terminal regions and low in the regions between them (Supporting Information 4). We expressed each Praja in SH-SY5Y cells and observed that Praja1 was expressed in both the cytoplasm and nucleus, whereas Praja2 and Anc-Praja were expressed only in the cytoplasm (Fig. 3B). This may be explained by the acquisition of a nuclear localization signal specific to Praja1 [2]. Notably, E3 ligases undergo auto-ubiquitination during their activity [19,20]. Anc-Praja was auto-ubiquitinated by MG132 treatment and possessed a RING domain sequence similar to Praja1/Praja2, suggesting that it also possesses E3 ligase activity (Fig. 3C). These results indicate that the reconstructed Anc-Praja can function as an E3 ligase, and its sequence and localization are more similar to those of Praja2 than to those of Praja1.

Anc-Praja and Tau were co-expressed in SH-SY5Y cells. However, no Anc-Praja-dependent reductions in Tau protein levels were observed (Fig. 4A, B). This suggests that Praja1 uniquely acquired the Tau recognition ability during its evolution. Therefore, we attempted to identify the evolutionary mutations that enabled the interaction of Praja1 with Tau. Praja1 underwent deletions in the N- and C-terminal regions after duplication from Praja2 (Fig. 4C). To verify whether these regions suppress the Tau recognition ability of Praja2, both regions were deleted from Praja2 (ΔN+ΔC-Praja2), and co-expressed with Tau. Interestingly, Tau was reduced in ΔN+C-Praja2 relative to wild-type Praja2, indicating that the N and C-terminal region disrupts Tau interaction with the Praja family (Fig. 4D). C-terminal region (682-707 aa) is a widely conserved 26 amino acid regions within the Praja2 sequences and is predicted to be an intrinsically disordered region, as shown by two disorder prediction programs (Fig. 4E) [21]. Moreover, two missense SNPs have been reported in humans (rs139484791: A705T and rs246105: A688D/V)[22-24]. Notably, rs246105 had a higher allele frequency in a group of non-small cell lung cancer patients with recurrence within three years after surgery, suggesting that the corresponding region plays a significant physiological role. These results suggest that the N and C-terminal regions of Praja2 may function as an auto-inhibitory region for interaction with Tau and that the loss in these regions might have enabled Praja1 to interact with Tau.

**Figure 4.**
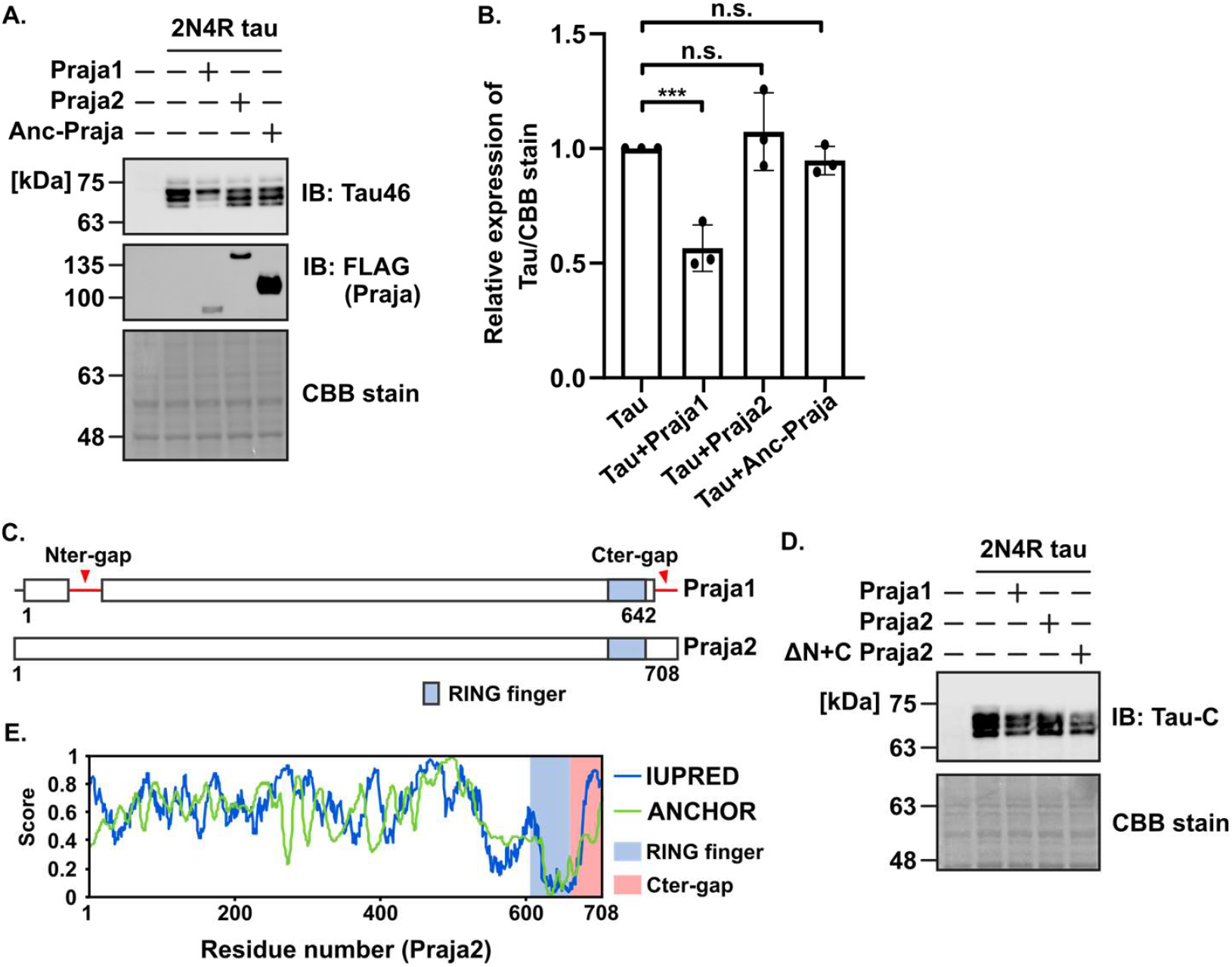
Praja1 may have started recognizing Tau as a novel substrate during its evolution. A. FLAG-Praja and Tau were co-expressed in SH-SY5Y cells and western blotted. B. Quantification of the results of A. C. Alignment of Praja1 and Praja2. Red arrows indicate regions lost during Praja1 evolution. Blue indicates the RING finger domain. D. Co-expression of ΔN+C Praja2 and Tau in SH-SY5Y cells and its western blot analysis. E. Analysis of disordered regions of Praja2 by IUPRED (blue line) and ANCHOR (green line). The blue region represents the RING finger and the red region represents the Cter-gap. For statistical analyses, mean values were subjected to multiple testing by ANOVA with post hoc Dunnett test (*p < 0.10, **p < 0.05, ***p < 0.01).

### 2.4 Praja1 interacts with FTDP-17-specific variant P301L Tau

To understand the pathological significance of Tau degradation by Praja1, we focused on the P301L mutant among the missense variants often found in patients with FTDP [11,25-27]. P301L Tau is known to be excessively phosphorylated, aggregated, and to interact abnormally with its binding partners [28,29]. Therefore, each Praja family member was tested for its effect on the P301L mutation. P301L Tau was transiently expressed in SH-SY5Y cells alone or together with Praja1 or Praja2. Praja1 reduced the expression of P301L Tau, whereas Praja2 had no effect (Fig. 5A, B). To determine whether P301L Tau was degraded by the E3 activity of Praja1, the cells were treated with MG132 for 6 h. Treatment with MG132 restored Tau expression, which was suppressed by Praja1 (Fig. 5C, D). Furthermore, Praja1 co-expressed with P301L Tau was localized to the cytoplasm of SH-SY5Y cells (Fig. 5E). Additionally, Anc-Praja did not degrade P301L Tau or WT Tau. These results suggest that P301L Tau and WT Tau are substrates of Praja1 (Fig. 5F, G). Together, these data show Praja1 potentially degrades P301L Tau similar to WT Tau.

**Figure 5.**
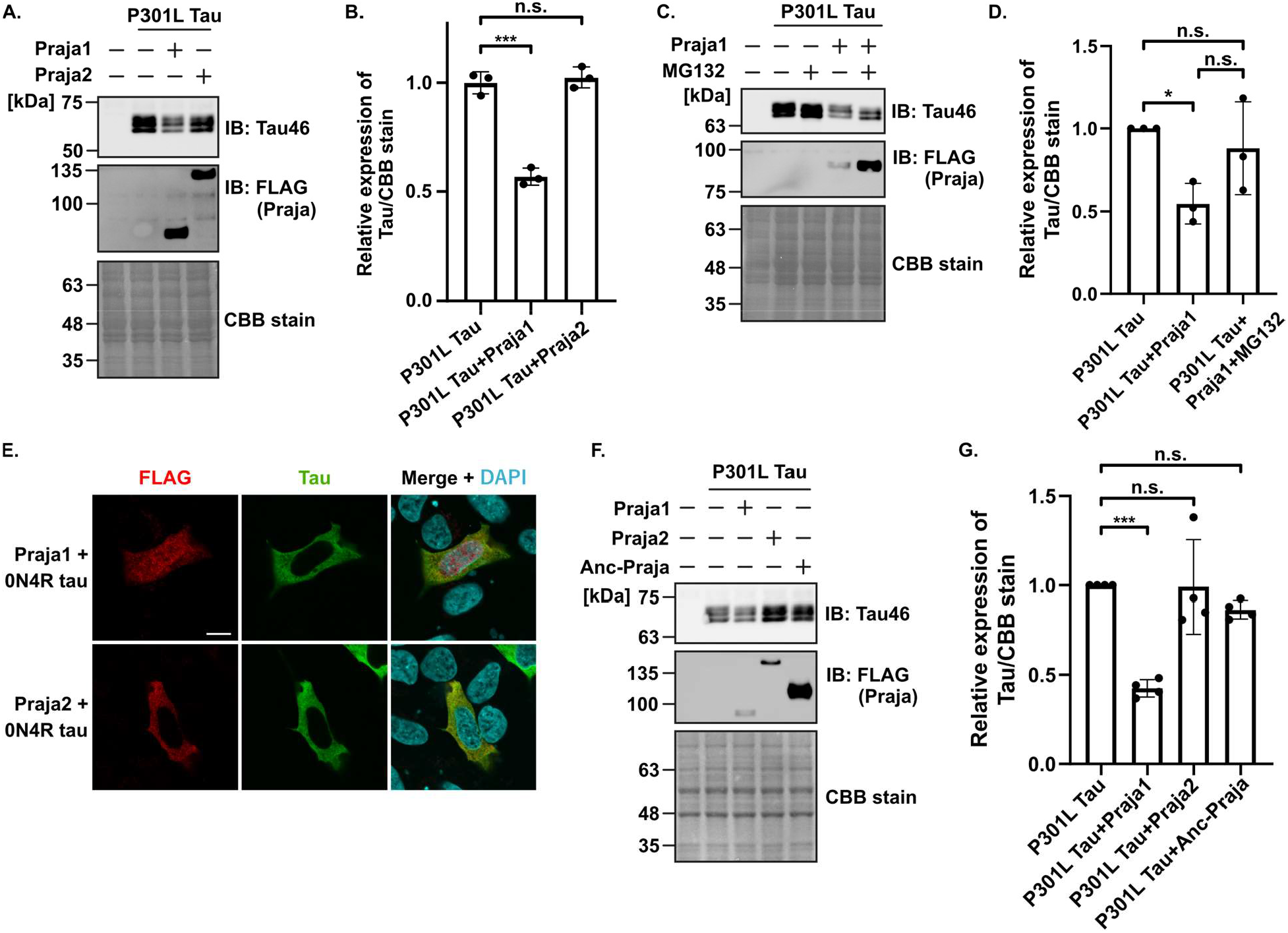
Praja1 recognizes P301L Tau in SH-SY5Y cells. A. FLAG-tagged Praja1/Praja2 and P301L Tau were co-expressed in SH-SY5Y cells and western blotted. B. Quantification of the results of A. C. FLAG-tagged Praja1 and P301L Tau were co-expressed alone or together in SH-SY5Y cells treated with or without MG132 for 6 h and western blotted. D. Quantification of the results of C. E. Immunostaining of FLAG-Praja and Tau in SH-SY5Y cells transfected with FLAG-tagged Praja1/Praja2 and P301L Tau alone or together. DAPI was used for nuclear staining. Scale bar: 10 µm. F. FLAG-Praja and Tau were co-expressed in SH-SY5Y cells and western blotted. G. Quantification of the results of F. For statistical analyses, mean values were subjected to multiple testing by ANOVA with post hoc Dunnett test (*p < 0.10, **p < 0.05, ***p < 0.01).

## 3. Discussion

In this study, we identified Tau as a potential substrate of Praja1 and discerned the regions evolutionarily involved in its interaction and relationship with the P301L Tau mutant. Notably, Tau was degraded only by the Praja1 paralog through its E3 ligase activity. Although direct binding was not observed between Praja1 and Tau using pull-down assay, BiFC-based fluorescent and Tau polyubiquitination induced by Praja1-Tau interaction was observed, indicating that Tau is a candidate substrate of Praja1. Further, Anc-Praja, the common ancestral sequence of Praja1 and Praja2, was reconstructed, and we showed that it could not degrade Tau, indicating that Praja1 uniquely acquired its Tau recognition ability during its evolution after the duplication event. Furthermore, the N and C-terminal regions lost in Praja1 in the common placental ancestor appears to be important for the recognition of Tau. Notably, this conserved region in Praja2 harbors SNPs that may be associated with disease risk in humans, suggesting that it is a functional region. Importantly, the disease-causing P301L Tau, which often interacts abnormally with its binding partners, was degraded by Praja1.

Here, Praja1 was shown to recognize Tau as a possible substrate protein. Tau hyperphosphorylation and aggregation are associated with several neurodegenerative diseases. This finding supports previous reports that Praja1 functions as a common suppressor of the aggregation of neurodegenerative disease-related proteins [7-9]. In contrast, Praja2, which has been previously reported to be downregulated in patients with AD, did not functionally interact with Tau at the protein level in this study [10]. This may be partially explained by the C-terminus of Praja2, which may interfere with its interaction with Tau. Not only the C-terminus but also the sequence identity between Praja1 and Praja2 diversified after gene duplication, especially at positions 100–500 of Praja2 [2].

The reason why Praja1 can bind to and inhibit the aggregation of multiple neurodegenerative disease-associated proteins, including tau, remains unresolved. This may be explained by the intrinsically disordered nature of Praja1, lacking a specific tertiary structure. E3 ligases containing disordered regions adopt suitable conformations in a partner-dependent manner [30,31]. In addition to the characteristics of Praja1, the biochemical properties of interacting proteins may also facilitate the recognition by Praja1. While other E3 ligases are sequestered from protein aggregates (e.g., polyQ and TDP-43 aggregates), Praja1 seems to successfully interact with aggregates for its clearance [7-9]. Further studies are needed to reveal the molecular mechanism by which Praja1 senses aggregated proteins and their pathological significance.

Tau is a disordered protein that flexibly adopts a specific conformation when interacting with microtubules [32,33]. Phosphorylation of Tau weakens its binding to microtubules, and free Tau may lead to pathological aggregation [34,35]. Most autosomal dominant FTLD-Tau mutations, including the P301L Tau mutation used in this study, occur in the microtubule-binding domain, resulting in accelerated phosphorylation of Tau owing to altered interactions with microtubules [11,24-26,36]. Additionally, the mutations may alter the interactions with Tau-binding proteins [32]. Despite this mutation, Praja1 promoted the degradation of Tau. Further investigation of the Praja1-Tau interaction is necessitated, as Tau has diverse proteoforms, including splice variants and post-translational modifications, perhaps using the E3 ligase CHIP, which specifically recognizes phosphorylated Tau [37]. It is also not possible to discuss the differences among the six isoforms of human Tau. Especially, as 2N4R Tau was used in this study, effect of N region deletion on interaction with Praja1 needs to be elucidated. In conclusion, our study on the regulation of Tau proteostasis by Praja1 may provide insights into the pathogenesis of tauopathy.

## 4. Methods

### 4.1 Plasmid construction

pCMV6-Entry plasmid vectors encoding FLAG-tagged human Praja1 and Praja2 were previously constructed [2]. Site-directed mutations were generated using a KOD-Plus-Mutagenesis Kit (#SMK-101, TOYOBO, Osaka, Japan). Each reaction was performed according to the manufacturer’s instructions. Plasmids were sequenced to confirm their DNA sequences. The following primers were used for each construct: C612A, H614A Praja1;

5’-GCCCACGCCTATTTCCACAAGCCGTGTG-3’

5’-CGGCAGCTCAGTTGCCACCTC-3’

ΔN-Praja2;

5’-AAAAGTGAAACAGAAATTCCCACTTG-3’

5’- AGCCCGACCTAAGCTTCTTTCAT-3’,

ΔC-Praja2;

5’-AAAAGTGAAACAGAAATTCCCACTTG-3’

5’- AGCCCGACCTAAGCTTCTTTCAT-3’,

P301L Tau;

5’-CGGGAGGCGGCAGTGTGCAAATAG-3’

5’-GGACGTGTTTGATATTATCC-3’.

### 4.2 Cell culture and gene transfection

SH-SY5Y cells were cultured in low-glucose Dulbecco’s modified Eagle’s medium (#041-29775, Wako Chemicals, Richmond, VA, USA) supplemented with 10% fetal bovine serum and 1% penicillin-streptomycin at 37 °C under 5% CO_2_. For proteasome inhibition, SH-SY5Y cells were incubated with 10 µM MG132 (#3175-v, Peptide Institute Inc., Osaka, Japan) dissolved in DMSO. Plasmid vectors were introduced using Polyethyleneimine Max (#24765-100, Polysciences, Warrington, PA, USA) or Lipofectamine 3000 (#L3000008, Thermo Fisher Scientific, Waltham. MA, USA), and experiments were conducted 16–24 h post-transfection.

### 4.3 Antibodies

Anti-DYKDDDDK tag (#66008-4-IG, Proteintech, Rosemont, IL, USA), HRP-conjugated anti-mouse IgG (H+L) (#SA00001-1, Proteintech), anti-Tau-C (#TIP-TAU-P04, Cosmo Bio, Tokyo, Japan), anti-Tau-360-380 (#TIP-TAU-P02, Cosmo Bio), and anti-Tau (#10274-1-AP, Proteintech) were used. Tau (Tau46) mouse monoclonal antibodies (4019) were purchased from Cell Signaling Technology, Danvers, MA, USA. Anti-DDDDK-tagged pAbs (anti-FLAG, PM020), anti-multi-ubiquitin mAbs (FK2, D058-3), and anti-His-tagged mAbs (OGHis) were purchased from MBL, Japan. Anti-mouse IgG (H+L) Alexa Fluor™ 488 conjugated (R37114), goat anti-rabbit IgG (H+L) cross-adsorbed secondary antibody, Alexa Fluor™ 594 (A11037) were purchased from Thermo Fisher Scientific. Phospho-Tau (Ser202 and Thr205) antibodies (AT8, MN1020) were purchased from Invitrogen, Waltham, MA, USA. Anti-FLAG-tag mAb (F1804-200UG) was purchased from Sigma-Aldrich, St. Louis, MO, USA.

### 4.4 Western blotting and immunoprecipitation

SH-SY5Y cells were lysed in cell lysis buffer (50 mM Tris-HCl (pH 7.8), 150 mM NaCl, 1% NP-40, and 0.5% sodium deoxycholate) supplemented with a complete protease inhibitor cocktail (#11836153001, Roche, Basel, Switzerland). The cell lysate was centrifuged at 15,000 rpm for 5 min at 4 °C, and the supernatant was treated with sample buffer solution (#198-13282, Wako Chemicals) and 3% 2-mercaptoethanol for 2 min at 95 °C. Protein concentrations were measured using a Pierce BCA Protein Assay Kit (#23225, Thermo Fisher Scientific) following the manufacturer’s protocol. For SDS-PAGE, Extra PAGE One Precast Gel 7.5% (#13070-74, Nacalai, Kyoto, Japan) was used, and the proteins were subsequently transferred to a PVDF membrane (#IPVH00010, Merck Millipore, Burlington, MA, USA). The membranes were washed with TBS-T (20 mM Tris (pH 7.5), 500 mM NaCl, and 1% Tween20) for 5 min and blocked with TBS-T+5% skim milk for 30 min at room temperature. The PVDF membrane was then incubated overnight with a primary antibody solution diluted in 2.5% skim milk and incubated for at least 1 h with an HRP-conjugated secondary antibody solution diluted in 2.5% skim milk. Then, the PVDF membrane was treated with the chemiluminescence reagent Immunostar LD (#292-69903, Wako Chemicals) for 10 s to 1 min, and images were acquired using a ChemiDoc Touch MP (Bio-Rad, Hercules, CA, USA). The same membrane was subjected to CBB staining (#SP-4011, APRO Science, Tokushima, Japan) for the quantitative evaluation of total proteins. The results were quantified using ImageJ and statistically analyzed using GraphPad Prism (GraphPad Prism 8 for Windows, GraphPad Software, Boston, MA, USA, www.graphpad.com).

For immunoprecipitation, proteins were extracted as described above, and protein G mag Sepharose (#28951379, Cytiva, Marlborough, MA, USA) conjugated with Tau-C antibody was used. Protein G MagSepharose and Tau-C antibodies were added and incubated at room temperature for 30 min. Protein extracts were then mixed at room temperature for 30 min, followed by the addition of sample buffer, and elution of bound proteins was performed at 95 °C for 5 min.

### 4.5 Immunofluorescence and microscopy

Immunofluorescence analysis was performed as previously described [2]. The SH-SY5Y cells were washed with PBS and fixed in 4% paraformaldehyde for 20 min at room temperature. The fixed cells were permeabilized with 0.3% Triton X-100 in PBS for 10 min at room temperature. The permeabilized cells were blocked with 5% BSA/PBS for 30 min at room temperature. Cells were then incubated overnight at 4 °C with the primary antibody dissolved in 2.5% BSA/PBS. Cells were incubated with the same solvent and secondary antibody for 2 h at 4 °C. Cell nuclei were stained with Vectashield mounting medium for fluorescence using DAPI (#H-1200, Vector Laboratories Inc., Newark, CA, USA). Images were captured using a confocal laser scanning microscope (#FV-3000, Olympus, Tokyo, Japan).

### 4.6 BiFC Assay

To investigate the protein-protein interactions between Tau and Praja1, we employed the BiFC assay []. The Venus fluorescent protein (Ex515/Em528) split into two fragments, VN-155 (I152L, 1-154 aa) and VC-155 (155-239 aa), were fused to Tau and Praja1 together with the linker sequence RSIAT. DNA sequences of Venus fragments with the linker sequence were synthesized in Eurofins Scientific, Louisville, KY, USA. VN-155 (I152L) was fused to the N-terminal of Praja1, and VC-155 was fused to either the N-terminal or C-terminal of Tau. We used N-terminally tagged Tau as C-terminally tagged Tau did not show Venus-derived fluorescent (data not shown). For the subcloning of Tau and Venus fragments into pCMV6-Entry vector, the HiFi Assembly Kit (NEB, #E2621) was utilized.

### 4.6 Construction of ancestral Praja1

A common ancestor sequence of Praja1 and Praja2 was reconstructed using 202 previously collected mammalian Praja family sequences [2]. GRASP was used to reconstruct the ancestral sequence using the joint reconstruction option [17]. Using this option, the most probable state in the combination of all ancestral states was calculated by optimizing the likelihood over the entire phylogenetic tree. The amino acid sequence obtained by GRASP was codon-optimized for *Homo sapiens*, and plasmid DNA was obtained by gene synthesis from Eurofins Scientific, Louisville, KY, USA.

### 4.7 Protein purification and pull-down Assay

The human Praja1 sequence was ordered from Eurofins Scientific with codon optimization in *E. Coli*. This sequence was inserted into the pET-HisTEV vector and transformed into Rosetta2 (DE3) cells (#71397, Merck Millipore) to express N-terminally His-tagged Praja1. Colonies picked from the above were pre-cultured in 100 mL of LB medium overnight and then grown in 2000 mL of medium for 1.5 h. When OD_600_ reached 0.6–0.8, His-Praja1 expression was induced with 0.5 mM Isopropyl-β-D-thiogalactopyranoside. The centrifuged pellet was resuspended in 1.5 mL of sonication buffer (50 mM Tris-HCl (pH 8.0), 150 mM NaCl, and a protease inhibitor) and subjected to ultrasonication. After centrifugation at 15,000 rpm, 10 min, at 4 °C, the supernatant was purified using a 0.45 µm PVDF filter and used as the input sample to be used for purification. Purification of this sample was performed using a HisTrap HP (5 mL; #17524802, Cytiva) on an Akta avant25 column (Cytiva). The column was equilibrated with 50 mM Tris-HCl (pH 8.0), 150 mM NaCl, and 50 mM imidazole, injected, and washed with the same buffer. For the elution of His-Praja1, 50 mM Tris-HCl (pH 8.0), 150 mM NaCl, and 500 mM imidazole were used. The fractions containing His-Praja1 were ultrafiltered, concentrated with 30 kDa NMWL Amicon Ultra (#UFC503024, Merck Millipore), and used for downstream analysis. We confirmed the presence of His-Praja1 in the eluted fraction using SDS-PAGE and western blotting with an anti-His tag. Additionally, 1 h of TEV protease treatment was used to verify whether the His-tag was removed from Praja1 and to confirm that the purification was successful.

A pull-down assay was performed using a Capturem His-Tagged Purification Miniprep Kit (#635710, Takara, Shiga, Japan). 50 µg of BSA was added to the column for blocking, followed by 5 µg of purified His-Praja1. The column was then centrifuged at 11,000 × *g* for 1 min. An extract of Tau-overexpressing SH-SY5Y cells was applied to the column and centrifuged under the same conditions. The extract was then eluted with wash and elution buffers containing 500 mM imidazole. The eluted fractions were treated with sample buffer and 2-Mercaptoethanol for 2 min at 95 °C and used as samples for western blotting.

## Abbreviations

ALS: amyotrophic lateral sclerosis
FTLD: frontotemporal lobar degeneration
AD: Alzheimer’s disease
PD: Parkinson’s disease
FTDP: frontotemporal dementia
BiFC: Bimolecular fluorescence complementation
GRASP: graphical representations of ancestral sequence predictions

## Author contributions

S.A., K.K., K.I., M.O., and W.O. performed the experiments. S.A. wrote the manuscript with support from W.O. and T.A., who supervised and designed the project. Funding was acquired by W.O. and T.A. Resources were provided by W.O. and T.A. All authors discussed the results and contributed to the final manuscript.

## Acknowledgments

This work was partially supported by the Global Consolidated Research Institute for Science, Wisdom, Waseda University and OPTORUN Co., Ltd. We would like to thank Editage (www.editage.jp) for the English language editing.

