## Supporting information for "E3 ligase Praja1 mediates ubiquitination and degradation of microtubule-associated protein Tau"

**Supporting information 1:** Western blot for FLAG-tagged Praja1/Praja2 expressed in SH-SY5Y cells.

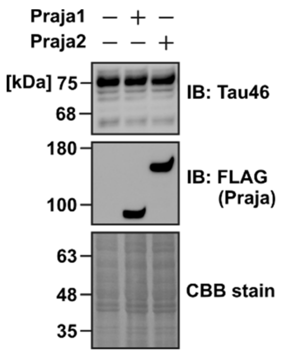

**Supporting information 2:** Purification of His-tagged Praja1 and pull-down assay with Tau. A. Chromatography by HisTrap HP to purify His-tagged Praja1 using AKTA Avant. Grey highlighted region was used for further experiment after ultrafiltered and concentrated with 30 kDa NMWL Amicon Ultra. B. CBB staining of purified His-Praja1 with TEV protease. Recognition site for TEV protease exist in between His-tag and Praja1. C. Western blot for purified His-tagged Praja1 with TEV protease treatment. D. Western blot for Tau from N-terminal His-tagged Praja1 purified from E. coli using pull-down assay.

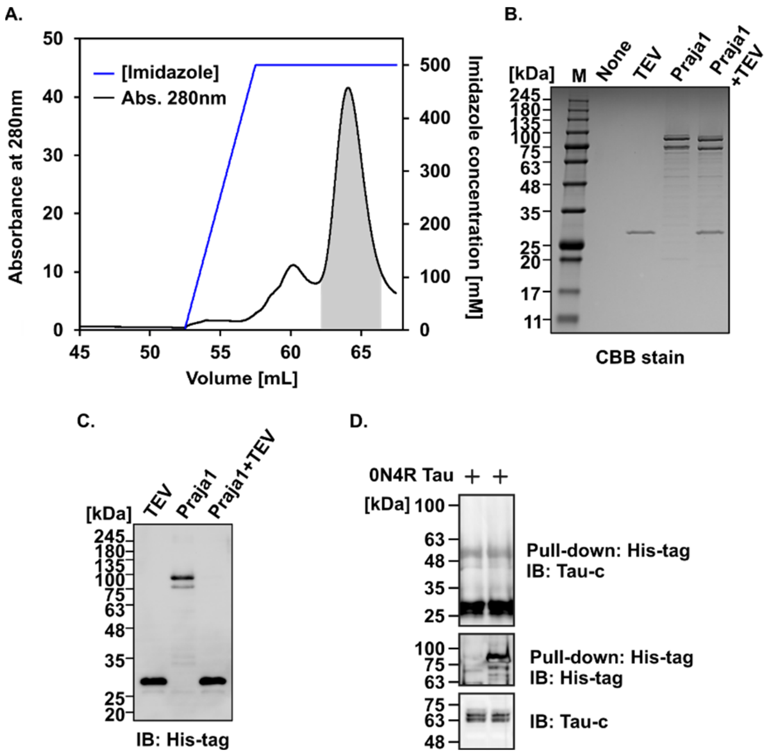

**Supporting information 3:** Multiple sequence alignment of human Praja1, Praja2 and ancestral Praja.

```
Anc_Praja                  -----------MGQESSKPTWPKPAGGYQTITGRRYGRRHAYVSFKPSMTRQERSSSQQS
Homo-sapiens_Praja2        MSQYTEKEPAAMDQESGKAVWPKPAGGYQTITGRRYGRRHAYVSFKPCMTRHERSLGRAG
Homo-sapiens_Praja1        -----------MGQESSKPVWPNPTGGYQSNTGRRYGRRHAYVSFRPPTSQRERIASQ--
                                      *.***.*..**:*:****: **************:*  :::**  .:

Anc_Praja                  GEYEGLELDDVQKEKASCCSPLDQVHPSLSGEPLLEKAGAEIPICRAALTQTTERSPSPF
Homo-sapiens_Praja2        DDYEVLELDDVPKENSSGSSPLDQVDSSLPSEPIFEKSETEIPTCGSALNQTTESSQS-F
Homo-sapiens_Praja1        -----------------------------------RKTNSEVPMHRSAPSQTTKRSRSPF
                                                              .*: :*:*   :* .***: * * *

Anc_Praja                  SAIHHSLEGSETSGSSLNLDNSSDSRYTPREYNASSVKNGIAFVHIDSYEPDGSDGEDNS
Homo-sapiens_Praja2        VAVHHSEEGRDTLGSSTNLHNHSEGEYIPGACSASSVQNGIALVHTDSYDPDGKHGEDND
Homo-sapiens_Praja1        STTRRSWDDSESSGTNLNIDNEDYSRYPPREYRASGSRRGMAYGHIDSYGADDSEEEGAG
                            : ::* :. :: *:. *:.* . ..* *    **. :.*:*  * *** .*... *. .

Anc_Praja                  QIELSAVKVETGKFQETLDDMLYDLEKG-VDSLSGLPPQFSSFNCEIREEFEELDPAPLV
Homo-sapiens_Praja2        HLQLSAEVVEGSRYQESLGNTVFELENREAEAYTGLSPPVPSFNCEVRDEFEELDSVPLV
Homo-sapiens_Praja1        PVERPPVRGKTGKFKDDK----LYDPEKGARSLAGPPPHFSSFSRDVREERDKLDPVPAA
                            :: ..   : .::::          :  . : :* .* ..**. ::*:* ::**..* .

Anc_Praja                  KYSSSDTEFVHQNNRTLQRSSAEDEVVAKKYQNNSSRERQRKNLTEDPARAPVDICNEQN
Homo-sapiens_Praja2        KSSAGDTEFVHQNSQEIQRS-SQDEMVSTKQQNNTSQERQTEHSPEDAACGPGHICSEQN
Homo-sapiens_Praja1        RCSASRADFLPQSSVASQSS---SEGKLATKGDSSERERREQNLPARPSRAPVSICG---
                           : *:. ::*: *..   * *   .*    .  :.:.:**: :: .  .: .*  **.

Anc_Praja                  TSDGEKNQGSSSELVVRPKVRKLASSSQVDQKTLFNSDEDK--QHSIERWRETSEAEENH
Homo-sapiens_Praja2        TNDREKNHGSSPEQVVRPKVRKLISSSQVDQETGFNRHEAK--QRSVQRWREALEVEESG
Homo-sapiens_Praja1        --GGENTSKSAEEPVVRPKIRNLASPNCVKPKIFFDTDDDDDMPHSTSRWRDTANDNEGH
                             . *:.  *: * *****:*:* *.. *. :  *: .: .   :* .***:: : :*.

Anc_Praja                  SDGLLR---KYDGENSSMFFDPRYS--EAQRETKNNQIKPETVTTQGRQAIVDNTFWNSC
Homo-sapiens_Praja2        SDDLLIKCEEYDGEHDCMFLDPPYSRVITQRETENNQMTSESGATAGR-QEVDNTFWNGC
Homo-sapiens_Praja1        SDGLAR---RGRGESSSGYPEPKYP--EDKREARSDQVKPEKVPRRRR-TMADPDFWTHS
                           **.*     .  ** .. : :* *.    :**:..:*:..*. .   *   .*  **. .

Anc_Praja                  EDYYRYYDKDEDSSECSDGEWSASLPHRFSGREKDQSSSDESWETLPGKEEHEPE-----
Homo-sapiens_Praja2        GDYYQLYDKDEDSSECSDGEWSASLPHRFSGTEKDQSSSDESWETLPGKDENEPE-----
Homo-sapiens_Praja1        DDYYKYCDEDSDS----DKEWIAALRRKYRSREQTLSSSGESWETLPGKEEREPPQAKVS
                            ***:  *:*.**    * ** *:* ::: . *:  ***.*********:*.**

Anc_Praja                  ----------------------LQSNSSGPEEENQELSLQEGEQTSLEEGEIPWLQYNEE
Homo-sapiens_Praja2        ----------------------LQSDSSGPEEENQELSLQEGEQTSLEEGEIPWLQYNEV
Homo-sapiens_Praja1        ASTGTSPGPGASASAGAGAGASAGSNGSNYLEEVREPSLQE-EQASLEEGEIPWLQYHE-
                                                   *:.*.  ** :* **** **:************:*

Anc_Praja                  NESSSDEENDPANEFVQPGVFMLDGNNNLEDDSSVSEDLDVDWSLFDGFADGLGVAEAIS
Homo-sapiens_Praja2        NESSSDEGNEPANEFAQP-AFMLDGNNNLEDDSSVSEDLDVDWSLFDGFADGLGVAEAIS
Homo-sapiens_Praja1        NDSSSEGDNDSGHELMQPGVFMLDGNNNLEDDSSVSEDLEVDWSLFDGFADGLGVAEAIS
                           *:***:  *:..:*: ** .*******************:********************

Anc_Praja                  YVDPQFLTYMALEERLAQAMETALAHLESLAVDVEVANPPASKESIDCLPETLVTEDHTA
Homo-sapiens_Praja2        YVDPQFLTYMALEERLAQAMETALAHLESLAVDVEVANPPASKESIDGLPETLVLEDHTA
Homo-sapiens_Praja1        YVDPQFLTYMALEERLAQAMETALAHLESLAVDVEVANPPASKESIDALPEILVTEDHGA
                           *********************************************** *** ** *** *

Anc_Praja                  VGQEQCCPICCSEYIKDEIATELPCHHFFHKPCVSIWLQKSGTCPVCRHVFPPALPEATA
Homo-sapiens_Praja2        IGQEQCCPICCSEYIKDDIATELPCHHFFHKPCVSIWLQKSGTCPVCRRHFPPAVIEASA
Homo-sapiens_Praja1        VGQEMCCPICCSEYVKGEVATELPCHHYFHKPCVSIWLQKSGTCPVCRCMFPPPL-----
                           :*** *********:*.::********:********************  ***.:

Anc_Praja                  ATSFLPDHDSPPSIHSATGTQ
Homo-sapiens_Praja2        APSSEPDPDAPPSNDSIAEAP
Homo-sapiens_Praja1        ---------------------
```
